# Evaluation of primer pairs for microbiome profiling across a food chain from soils to humans within the One Health framework

**DOI:** 10.1101/843144

**Authors:** Wasimuddin, Klaus Schlaeppi, Francesca Ronchi, Stephen L Leib, Matthias Erb, Alban Ramette

**Affiliations:** Institute for Infectious Diseases, University of Bern, Bern, Switzerland; Institute of Plant Sciences, University of Bern, Bern, Switzerland; Department for Biomedical Research, University of Bern, Inselspital, 3008 Bern, Switzerland

**Keywords:** One Health, microbiome, 16S rRNA gene, high throughput sequencing, soil, plant root, cow rumen, human gut

## Abstract

The “One Health” framework emphasizes the ecological relationships between soil, plant, animal and human health. Microbiomes play important roles in these relationships, as they modify the health and performance of the different compartments and influence the transfer of energy, matter and chemicals between them. Standardized methods to characterize microbiomes along food chains are, however, currently lacking. To address this methodological gap, we evaluated the performance of DNA extractions kits and commonly recommended primer pairs targeting different hypervariable regions (V3-V4, V4, V5-V6, V5-V6-V7) of the 16S rRNA gene, on microbiome samples along a model food chain, including soils, maize roots, cattle rumen, and cattle and human faeces. We also included faeces from gnotobiotic mice colonized with defined bacterial taxa and mock communities to confirm the robustness of our molecular and bioinformatic approaches on these defined low microbial diversity samples. Based on Amplicon Sequence Variants, the primer pair 515F-806R led to the highest estimates of species richness and diversity in all sample types and offered maximum diversity coverage of reference databases in *in silico* primer analysis. The influence of the DNA extraction kits was negligible compared to the influence of the choice of primer pairs. Comparing microbiomes using 515F-806R revealed that soil and root samples have the highest estimates of species richness and inter-sample variation. Species richness decreased gradually along the food chain, with the lowest richness observed in human faeces. Primer pair choice directly influenced the estimation of community changes (beta diversity) within and across compartments and may give rise to preferential detection of specific taxa. This work demonstrates why a standardized approach is necessary to analyse microbiomes within and between source compartments along food chains in the context of the One Health framework.

## Introduction

The “One Health” concept emphasizes the ecological relationships and interdependencies between humans, plants, animals and environmental health (Destoumieux-Garzon et al., 2018). Until recently, the One Health concept primarily focused on the origin and transfer of zoonotic pathogens, vectors of pathogens and antibiotic resistance between interacting entities (Destoumieux-Garzon et al., 2018). During the past decade, however, microbial communities (or microbiomes) have been shown to play important roles in connecting the humans, plants, animals and environment (van Bruggen et al., 2019). Thus, recommendations have been made to extend the One Health concept to include the full breadth of microbes (Bell et al., 2018;Trinh et al., 2018;van Bruggen et al., 2019). Adopting a microbiome perspective may strengthen the One Health concept due to i) the vital services provided by microbiomes to overall ecosystem health, ii) the importance of microbiome processes for the transfer of energy, matter and chemicals between compartments along the food chain, and iii) the important contribution of microbiomes to the health of the different hosts and compartments. However, methodological challenges remain to adequately characterize and allow comparison of the different microbiomes in order to track microbial transfer and to quantify the role of microbiomes in food chain health (Trinh et al., 2018).

Widely established approaches enable quantifying the diversity and richness of microbiomes with high resolution from diverse source compartments by sequencing the 16S rRNA marker gene amplified by ‘universal’ primers (Fricker et al., 2019). In order to meet the sequence length requirement of short-read sequencing technologies, various primer pairs have been designed to amplify short hypervariable regions of the 16S rRNA gene. Both, choice of hypervariable region of 16S and primer pair, influence the description of microbial diversity (Claesson et al., 2010). Thus, care should be taken in choosing appropriate primer pairs, as limited taxa coverage, over- or underrepresentation of taxa in a specific environment due to biases in primer amplification could produce unreliable results (Claesson et al., 2010). Free-living microbial communities such as those in soil and lake sediment may exhibit higher microbial richness than host-associated communities such as animal gut microbiomes (Thompson et al., 2017). Furthermore, among host associated communities, plant roots show higher microbial richness than other plant and animal associated microbial communities (Thompson et al., 2017). Because each source compartment is unique in terms of microbial richness and composition (Thompson et al., 2017), in studies where microbiomes from different source compartments are investigated, such as within the One Health framework, investigators should carefully select primers to avoid methodological biases and to maximize the detection of taxa (Trinh et al., 2018). To our knowledge, no study has systematically included diverse samples from different compartments in a primer comparison experiment. Thus, the key information about choice of primer pairs required to conduct One Health experiments is missing.

The prokaryotic primer pair 515F-806R, which was designed to detect both archaea and bacteria by amplifying V4 region, is recommended by the Earth Microbiome Project and has been extensively used to study soil microbiomes (Apprill et al., 2015;Parada et al., 2016;Walters et al., 2016). Nevertheless, two recent studies that evaluated the best performing primer pair based on taxa diversity coverage (Klindworth et al., 2013;Thijs et al., 2017), recommend the primer pair 341F-805R, which amplifies the V3-V4 region, over other primer pairs. In the *Klindworth et al.* study, 512 primer pairs were tested *in silico* against the SILVA v108 database (376,437 sequences) for amplification of archaeal and bacterial sequences. Yet, since that study 318,734 additional sequences have been added to the latest SILVA release v132, which almost doubled the size of the SILVA database. Hence, the previously characterized primer coverages should be re-examined using the enhanced current database. The study by *Thijs et al*. used both *in silico*, as well laboratory experiments, to access the best primer pair but did not include the 515F-806R primer pair in their comparisons and performed rather shallow sequencing (454 pyrosequencing) of soil samples. Several studies target exclusively bacteria to answer specific questions (Mazmanian et al., 2005;Hebbandi Nanjundappa et al., 2017), thus bacterial specific primer pairs should be included as well in such comparisons. Two primer pairs; 799F-1193R (V5-V6-V7) and 787F-1073R (V5-V6) have been preferably used in compartment specific studies, primer pair 799F-1193R in plant system due to reduced amplification of plant organelle DNA (Beckers et al. 2016) and 787F-1073R in mouse studies with limited diversity microbiomes (Li et al., 2015;Hebbandi Nanjundappa et al., 2017). Furthermore, the Divisive Amplicon Denoising Algorithm 2 (*DADA2*) introduced a model-based approach for identifying sequencing errors without the need of constructing OTUs and at the same time for detecting less false positives in comparison to earlier methods (Callahan et al., 2016). In light of the advancement in high-throughput sequencing methods and in high-resolution analysis methods, the choice of primer pairs should be re-examined in order to achieve higher taxonomic coverage.

In this study, our objective was to evaluate the performance of four commonly used primer pairs 787F-1073R, 799F-1193R, 515F-806R and 341F-805R for 16S amplicon sequencing especially within a One Health framework by including microbial communities from different source compartments along the human food chain; including samples from soil, plant, mouse, cattle and humans. We then used the best performing method to gain first insights into the commonalities and differences between the microbiomes along a model food chain.

## Material and Methods

### Sample collection

Samples were collected from four different source compartments with the aim to maximize the heterogeneity within compartment in the experiment (Supplementary Table 1). Briefly, five soil samples, each from a different soil type, six maize root samples from three different geographical locations, five cow samples including three faeces and two rumen samples, six human faeces samples from volunteers belonging to two couples, one child (3 years of age) and one female sample, were collected. Additionally, faeces from a gnotobiotic mouse strain colonized with defined microbial community (four bacteria of the Altered Schaedler’s Flora here referred as ASF.4: *Lactobacillus_acidophilus*_ASF360, *Lactobacillus_murinus*_ASF361, *Clostridium_sp*_ASF500, *Bacteroides_distasonis*_ASF519) and a mock microbial community DNA (8 bacterial + 2 yeasts species mixed in defined proportions) (ZymoBIOMICS Microbial Community DNA standard, Zymo Research, USA) were included in the experiments. All samples were stored at −80°C until further analysis.

### Bacterial DNA extraction and 16S rRNA gene amplicon sequencing

Samples were homogenized by bead beating at 50Hz for four minutes using a TissueLyser LT (QIAGEN, Germany). Genomic bacterial DNA was extracted from all samples by using DNeasy PowerSoil Pro kit (QIAGEN, Germany) according to the manufacturer’s instructions. As samples from diverse source compartments were included in the planned experiment, we additionally extracted bacterial DNA using kits, which are generally used for the particular source compartment, in order to examine the DNA extraction kit effect on source compartment microbiome (Knauth et al., 2013;Lim et al., 2018). We extracted soil and root samples with the NucleoSpin Soil DNA extraction kit (Macherey-Nagel, Germany) according to the manufacturer’s instructions (Knauth et al., 2013). Likewise, mouse faecal samples were extracted with the QIAamp DNA FAST Stool Mini Kit (QIAGEN, Germany) (Lim et al., 2018) following manufacturer’s instructions, however an additional step of lysozyme treatment was added as reported previously (Mamantopoulos et al., 2017).

Four primer pairs, namely 787F-1073R, 799F-1193R, 515F-806R and 341F-805R, were used to amplify the V5-V6, V5-V6-V7, V4, V3-V4 hypervariable regions of the 16S rRNA gene, respectively (Table1). Forward primers and reverse primers carried overhang adapters (5’ TCGTCGGCAGCGTCAGATGTGTATAAGAGACAG-Forward primer, 5’ GTCTCGTGGGCTCGGAGATGTGTATAAGAGACAG-Reverse primer) for compatibility with Illumina index and sequencing adapters. A two-round amplification process was used to amplify the DNA samples, while reducing dimer formation, which is often the problem in multi-primer, multi-template PCR, especially with primers containing long overhang regions (Kalle et al., 2014). Amplicon PCR reactions were carried out using the Faststart PCR system (Roche, Switzerland). The 25-μl PCR mix was composed of 3 ng/μl DNA, 1× FastStart PCR grade nucleotide mix buffer without MgCl_2_, 4.5 nM MgCl_2_, 200 μM each of PCR grade nucleotides, 0.05 U/μl Fast Start Taq DNA Polymerase, 400 nM target-specific primers, 5% DMSO and 9 μl of PCR certified water. PCR cycling conditions consisted of an initial activation step at 95° C for 3 min, followed by 32 cycles with denaturation at 95° C for 30 s, annealing at 62° C for 30 s, extension at 72° C for 30 s and final extension at 72° C for 10 min. PCR products were subsequently purified using SPRI based size selection (Beckman Coulter Genomics, USA) and quantified using Qubit 2.0 Fluorometer. Equal amount of first round purified PCR products were used as templates for the second round indexing PCR using Nextera XT Index kit (Illumina USA). Briefly, 50 μl of reaction mix consisted of 5 μl of first round PCR product (2.5 ng/μl), 5 μl of Nextera XT Index Primer 1, 5 μl of Nextera XT Index Primer 2, 25 μl of MyFi Mix (2x) (Bioline, Meridian Bioscience, France) and 10 μl of PCR certified water. Indexing PCRs cycling conditions were according to standard Illumina 16S metagenomic sequencing library preparation protocol (https://support.illumina.com/). Second-round amplicon libraries were purified using SPRI based size selection (Beckman Coulter Genomics, USA) and quantified using Fragment Analyzer (Agilent, USA). The final pooled libraries were paired-end sequenced (2×300 cycles) in a single run on Illumina MiSeq at the NGS platform of University of Bern (www.ngs.unibe.ch). Negative controls were included in both the DNA extraction (no DNA template added) and 16S PCR amplification (with PCR certified water) to test for contamination. No noticeable DNA contamination of the negative controls after PCR amplification was observed during quantification using Qubit 2.0 fluorometer and by Fragment Analyzer.

**Table 1.**
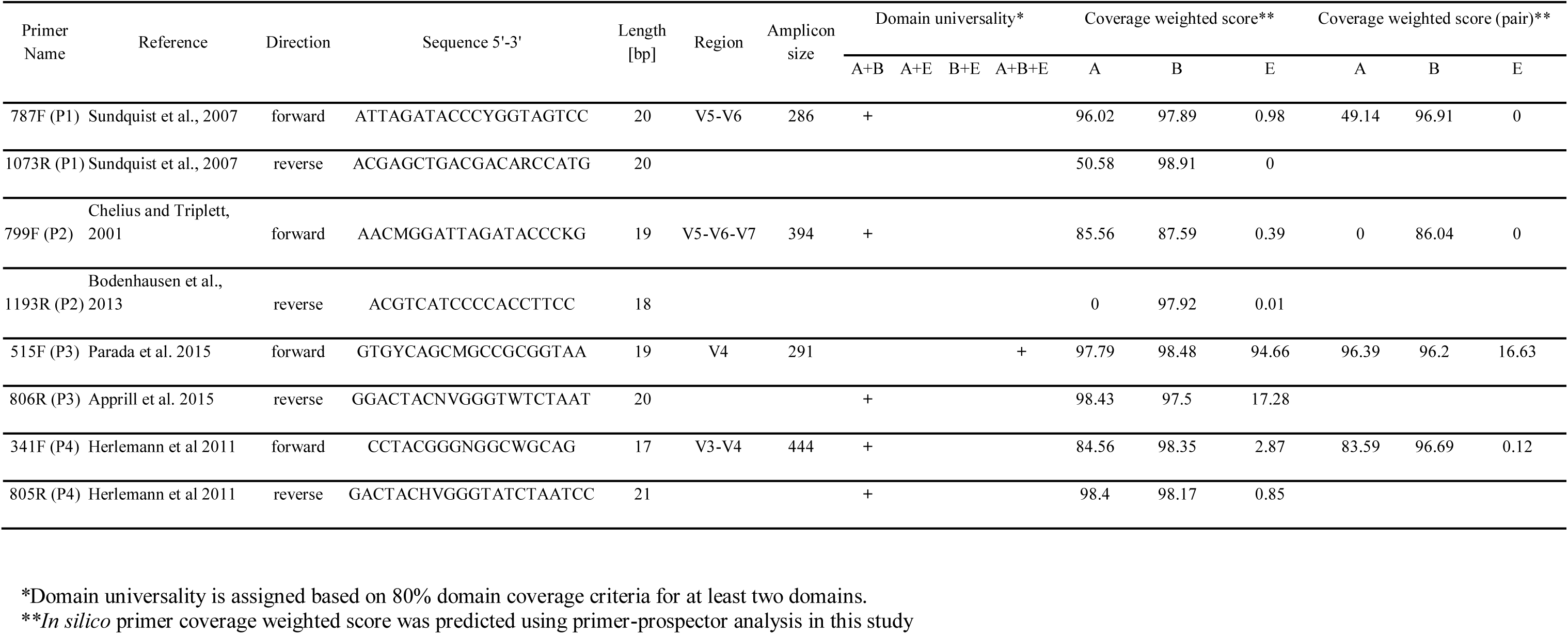
Details of the primers used in the current study and their *in silico* evaluation.

### Bioinformatics

Demultiplexed reads without barcodes and adapters were received as output from the sequencing centre. All subsequent analyses were performed within the R environment (R version 3.5.1, R Development Core Team 2011). For data pre-processing, we followed the *DADA2* pipeline (version 1.10; Callahan et al., 2016) by adjusting parameters to each of the four primer pair datasets. For each primer pair dataset, reads were trimmed from both ends based on quality profile, error rates were learned from the data using the parametric error model as implemented in *DADA2*. After denoising and merging, chimeric sequences (bimera) were removed from the datasets by following the ‘consensus’ method as implemented in *DADA2.* The final table thus consisted of a tabulation of number of occurrences of non-chimeric amplicon sequence variants (ASVs; i.e. sequence differing by as little as one nucleotide) in each sample. Taxonomy assignments of representative ASVs were performed using the naïve Bayesian classifier method with the latest SILVA v132 non-redundant (NR) database. SILVA database was chosen because it also contains eukaryotic sequences, which would be helpful to determine whether the primer pairs also amplify eukaryotic sequences. Species level assignment was done by exact matching (100% identity) of ASVs with database sequences, as previously recommended (Edgar, 2018). *Phyloseq* (version 1.24.2, McMurdie and Holmes, 2013) package was used for further data processing, and ASVs belonging to chloroplast, mitochondria, and unassigned ASVs at phylum level were removed from the dataset. We merged ASV read abundance profiles based on their phylum, genus and species level assignments to analyse the microbiome diversity across the four datasets produced by the different primer sets.

### Alpha, beta diversity analysis and differential species abundance

We investigated the effects of DNA extraction kits, primer pair, and source compartment on microbial diversity for each sample by using two different alpha diversity indices (number of observed species and Shannon) after rarefying the data to 10,300 sequences per sample using *Phyloseq*. To analyse the association of DNA extraction kits, primer pair and source compartment with these alpha diversity matrices, we performed General Linear Modelling (GLM) by using the *lme*_*4*_ package in R (Bates et al., 2015). We included primer pairs (787F-1073R, n= 48; 799F-1193R, n= 46; 515F-806R, n= 46 and 341F-805R, n= 47), source compartment (mock; n= 20, mouse; n= 39, soil; n= 38, root; n= 48, cow; n= 20 and human; n= 22), DNA extraction kit (source specific kits; n=61, Powersoil; n=126) and interaction between primer pair and source compartment (primer pair*source compartment) in the model as explanatory variables for each alpha diversity metric table.

Beta diversity analyses were based on calculated Jaccard and Bray-Curtis dissimilarity matrices after rarefying the data to 10,300 sequences per sample by using *Phyloseq*. The permutational multivariate analysis of variance (PERMANOVA) was employed as implemented in the *adonis* function of the *vegan* package (version 2.5-2; Oksanen et al., 2019) to test the significance of the differences in community composition with 999 permutations. For both beta diversity metrics, we similarly included DNA extraction kit, primer pairs, source compartment, and interaction between primer pair and source compartment (primer pair*source compartment) in the models as explanatory variables. To visualize patterns of separation between different sample categories, non-metric multidimensional scaling (NMDS) plots were prepared based on the Bray-Curtis dissimilarity coefficient. To understand whether choice of primer pair reflects true shift in microbial community composition or differential spread (dispersion) of data points from their group centroid, we assessed the multivariate homogeneity of group dispersions (variances) using the *betadisper* function of the *vegan* package.

In order to identify the species accountable for differences in grouping by primer pair, we employed a negative binomial model-based approach available in the *DESeq2* package in R (Love et al., 2014). Wald tests were performed and only species remaining significant (p<0.01) after the Benjamini–Hochberg correction were retained.

### *In silico* primer analysis

We estimated the primer pair’s predicted coverage and mismatches to the target database with *PrimerProspector* (Walters et al., 2011). For this purpose, we used the latest SILVA v132 NR 16S rRNA gene database with 695,171 sequences. Primers weighted scores were calculated with PrimerProspector’s default formula. Additional penalty score of 3.00 was given, if the final 3’ base of primer had a mismatch with its target sequence. Lower value of weighted score suggests better primer performance, whereas values above 0 suggest poor performance of primer pairs (Walters et al., 2011). Predicted coverage of the primer pairs was calculated at domain and phylum levels. We attributed domain universality to a primer sequence at a stringency criterion of 80%, i.e. only when a primer sequence showed 80% or more coverage of at least two taxonomic domains.

## Results

### Read output and taxa distribution

The Illumina MiSeq sequencing of different regions of 16S rRNA gene amplified by four primer pairs generated roughly 17 million raw reads in total with on average 84,883 reads per sample. Total number of raw reads differed according to primer pair, with highest number of reads associated with 787F-1073R (5,408,844 reads) followed by 515F-806R (4,898,357 reads), 799F-1193R (3,687,401 reads) and 341F-805R (3,151,874 reads). Different proportions of reads were filtered out at each step of quality filtering, denoising, merging and chimera removing for each primer pair (Supplementary Table 2) and in the end, the highest number of reads was retained for 515F-806R (72.6%), followed by 799F-1193R (51.8%), 341F-805R (49.5%), and 787F-1073R (37.7%). In terms of read taxonomic classification at the domain level, no reads were found to belong to Eukaryota for all primer pairs, however, distribution of reads assigned to Bacteria and Archaea and for Chloroplast and Mitochondria differed according to primer pair used (Supplementary Table 3). From four diverse source compartments together with the two low diversity controls (mouse and mock bacterial community), in total 43 different phyla were observed after removing chloroplast and mitochondria sequences with half (n=21) of the phyla commonly identified by all primer pairs (Figure 1a). The highest number of phyla (n=42) was detected by the primer pair 515F-806R; yet the latter did not detect the Caldiserica phylum, which was only detected by 341F-805R in a single soil sample S2B with only two reads. Identified phyla differed in terms of relative abundance across source compartments but also among primer pairs within a compartment (Figure 1b). As most of the ASVs were not assigned at the species level, the genus level was chosen as a common taxonomic level for dataset comparison across primer pairs. At genus level, a total of 955 unique genera were identified, with only 348 commonly identified by all four primer pairs (Figure 1c) and the largest number (n=696) identified by primer pair 515F-806R.

**Figure 1.**
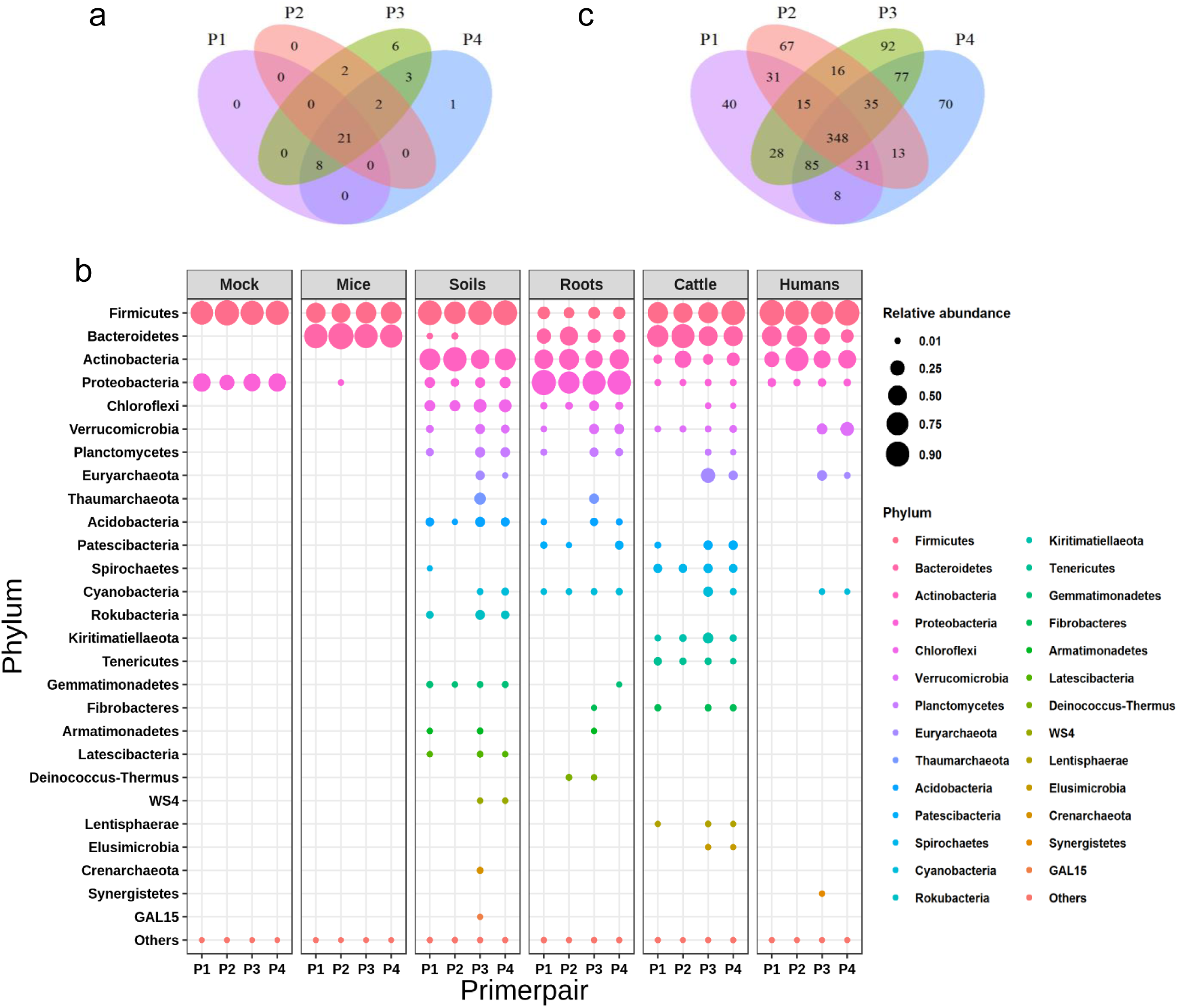
Comparative evaluation of primer pairs based on number and relative abundance of assigned taxa. Number of taxa shared and uniquely identified by primer pairs (787F-1073R (P1), 799F-1193R (P2), 515F-806R (P3) and 341F-805R (P4)) at **(a.)** phylum and **(c.)** genus level. Out of total 43 different phyla and 955 genera, 21 phyla and 348 genera are commonly identified by all four primer pairs. **(b.)** Bubble plot showing relative abundance of phylum by each primer pair and across source compartments. Size of the circles are proportional to relative abundance of phylum. Phyla are arranged according to total decreasing relative abundance and rarer phyla (<0.01%) are jointly included as ‘Others’.

With the mock community samples, we observed consistent performance of all primer pairs: out of eight bacterial species with negligible impurity (<0.01% foreign microbial DNA according to supplier), all eight bacterial species were recovered, however maximally assigned at genus level (Supplementary Figure 1). Only with primer pair 787F-1073R and 799F-1193R, we observed an additional taxon *Parabacteroids*, which was present in few (five) samples with <20 reads, which could be due to contamination or impurity. Removing rare ASVs from the dataset is recommended based on the number of samples in which they are present or by total count, because *DADA2* may be more sensitive to low amount of contamination (Caruso et al., 2019). However, we avoided using such filtering of rarer ASVs as in our dataset each sample is almost unique in terms of microbiome (due to the choice of heterogeneous samples in each compartment) and each primer pair dataset can be differentially influenced by such filtering parameter. Most reads from all primer pairs were successfully assigned at genus level for the mock community, however small proportions of taxonomically unassigned ASVs (at genus level) were present (Supplementary Figure 1). With respect to genus relative abundances, the primer pair 799F-1193R was noted to be biased towards *Bacillus* with more than half of reads assigned to this genus, and thus detected relatively fewer reads of all other genera (Supplementary Figure 1).

### Choice of primer pair influences alpha and beta diversity

We used General Linear Modelling (GLM) to test whether differences in alpha diversity estimates (either number of observed species, or Shannon) between samples could be explained by DNA extraction kit, primer pair, source compartment and the interaction between primer pair and source compartment (primer pair*source compartment). Significant effects were observed for primer pair (observed number of species, p<0.001; Shannon, p<0.001) and source compartment (observed number of species, p<0.001; Shannon, p<0.001; Supplementary Table 4) on tested alpha diversity indices. However, no effect of DNA extraction kit (observed number of species; p=0.846, Shannon; p=0.188) was observed on both tested alpha diversity indices (Supplementary Table 4, Figure 2). Interaction between primer-pair and source compartment showed no significant effect on observed species (p=0.171), but marginal effect on Shannon (p=0.042; Supplementary Table 4, Figure 2). Out of the four diverse source compartments, soil showed the highest microbial diversity, while the mouse and mock community samples (with defined bacterial strains) as expected showed the lowest (Figure 2). Similarly, primer pair 515F-806R revealed highest microbial diversity and 799F-1193R lowest (Figure 2).

**Figure 2.**
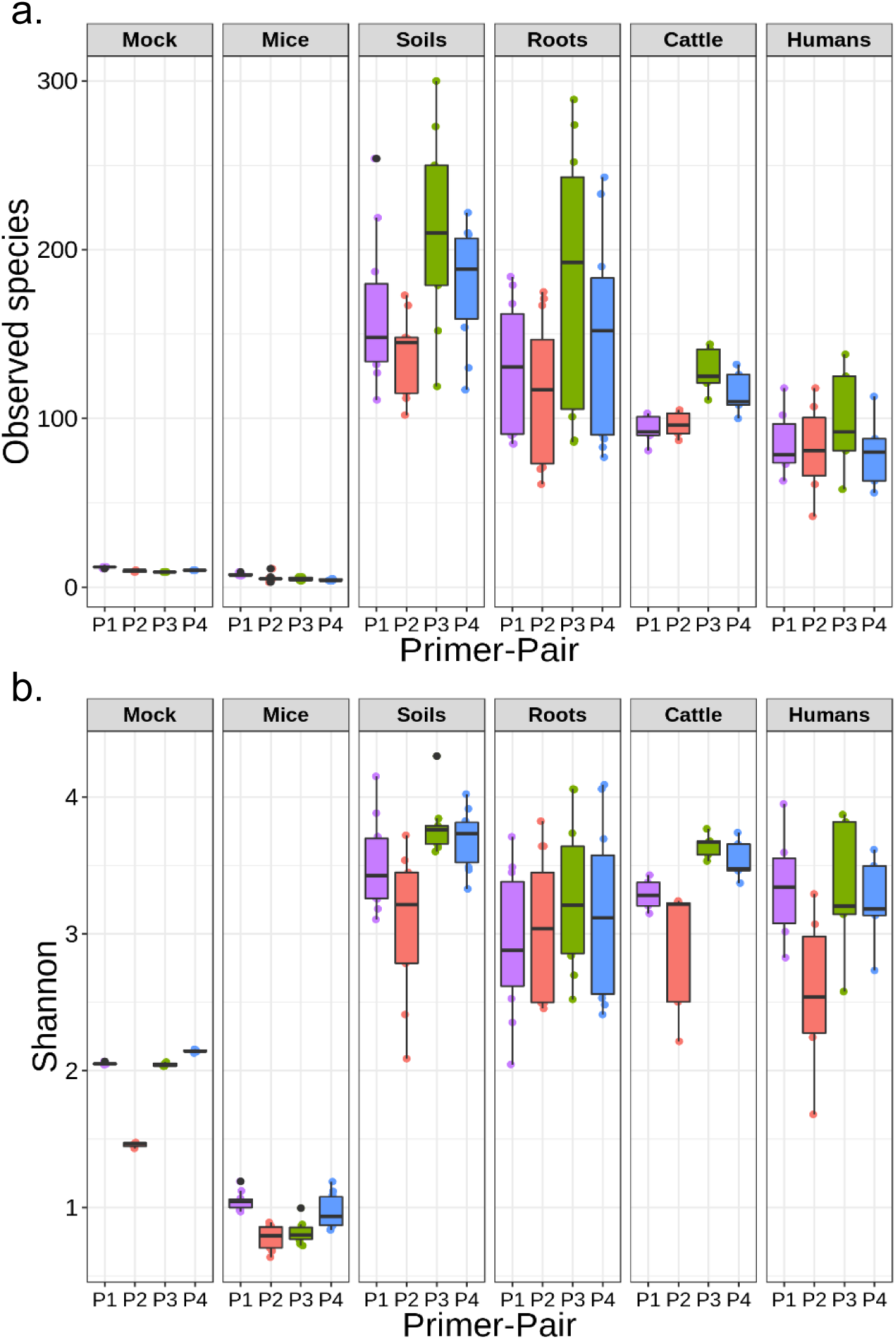
Box plots showing microbial alpha diversity of source compartments revealed by each primer pair. Difference in **(a.)** number of observed species and **(b.)** Shannon diversity of each source compartment by the primer pairs used (787F-1073R (P1), 799F-1193R (P2), 515F-806R (P3) and 341F-805R (P4)). For most source compartments, primer pair 515F-806R (P3) reflected the highest microbial diversity and 799F-1193R (P2) the lowest. Soil samples showed the highest alpha diversity and gnotobiotic mouse samples with defined colonized bacteria the least.

To determine whether choice of primer pair influences microbial community composition, we calculated two beta diversity metrics (Jaccard and Bray-Curtis) and included again DNA extraction kit, primer pair, source compartment and the interaction primer pair*source compartment as explanatory variables in PERMANOVA models. We observed a significant effect of primer pair (Jaccard: R^2^=0.024, p=0.001; Bray-Curtis: R^2^=0.021, p=0.001) and source compartment (Jaccard: R^2^=0.486, p=0.001; Bray-Curtis: R^2^=0.624, p=0.001), but also interaction primer pair*source compartment (Jaccard: R^2^=0.106, p=0.001; Bray-Curtis: R^2^=0.081, p=0.001) on microbial beta diversity estimates (Figure 3, Supplementary Table 5). However, no effect of the DNA extraction kit (Jaccard: R^2^=0.003, p=0.052; Bray-Curtis: R^2^=0.002, p=0.142) on the microbial community composition was observed (Supplementary Table 5). Homogeneity of dispersion analysis for primer pairs using *betadisper* function suggested true homogeneity in dispersions (Jaccard: p=0.831; Bray-Curtis: p=0.616; Supplementary Figure 2).

**Figure 3.**
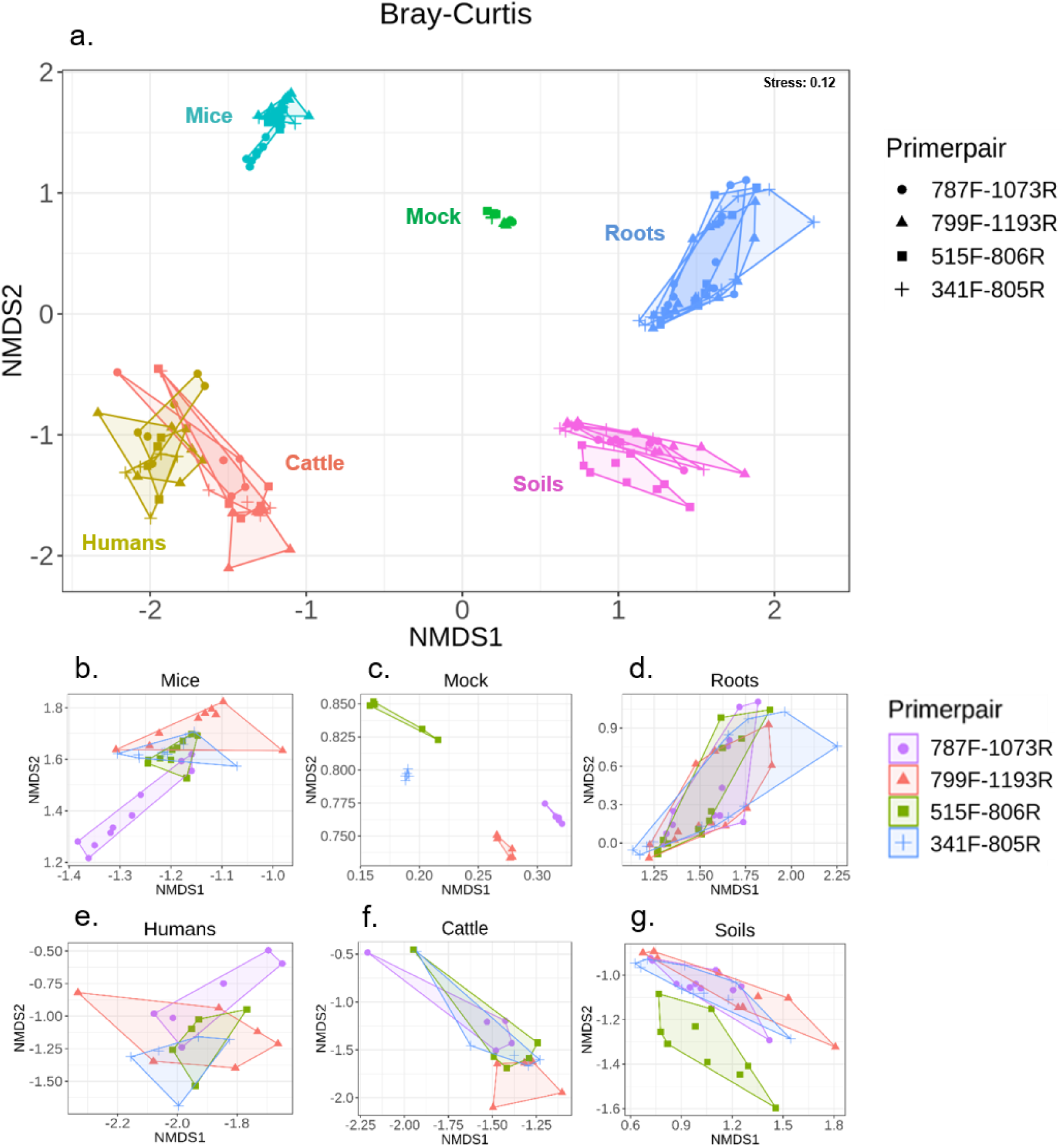
NMDS plots showing microbial beta diversity of source compartments based on Bray-Curtis distance metric. NMDS plots reflecting differences in microbial community composition of **(a.)** all source compartments together and for each source compartment (extracted values from the main NMDS plot); **(b.)** mice, **(c.)** mock, **(d.)** roots, **(e.)** humans, **(f.)** cattle, **(g.)** soils based on Bray-Curtis distance metric calculated using four primer pairs.

### Primer pairs are associated with differential abundance of bacterial species

To determine whether the choice of primer pair influences the ability to detect differences in relative species abundance, we employed negative binomial-based Wald tests. We compared species abundances obtained with 515F-806R, as it detected highest number of species, against the species abundances obtained with all other primer pairs (787F-1073R, 799F-1193R, 341F-805R). We identified 58 bacterial species that differed significantly (p<0.01) in abundance between primer pairs 515F-806R and 787F-1073R, with 19 (32.8%) and 39 (67.2%) showing significant decrease and increase in mean relative abundance, respectively (Figure 4a). Highest number of differential abundant species belonged to the phylum Firmicutes (19 species) followed by Actinobacteria (14 species). Five members of Archaea (3 species; Thaumarchaeota, 2 species; Euryarchaeota) showed higher abundance with 515F-806R than with 787F-1073R (Figure 4a). Analysing the primer pair 515F-806R against 799F-1193R revealed 86 bacterial species, significantly different in abundance, with 19 (22.1%) and 67 (77.9%) species showed significant decrease and increase in mean relative abundance, respectively (Figure 4b). Bacterial species showing differential abundance were mainly from the phylum Firmicutes (21 species), followed by Actinobacteria (18 species). Similar to comparison between 515F-806R and 787F-1073R, five members of Archaea (3 species, Thaumarchaeota; 2 species, Euryarchaeota) showed higher abundance with 515F-806R than with 799F-1193R (Figure 4b). Likewise, 25 bacterial species showed significant difference in abundance between primer pair 515F-806R and 341F-805R, whereas 9 (36.0%) and 16 (64.0%) showed significant decrease and increase in mean relative abundance, respectively (Figure 4c). Similar to previous comparisons, highest number of bacterial species showing differential abundance were found for Firmicutes (11 species) and archaeal members (3 species) were higher in abundance with 515F-806R. Overall, among all the comparisons, primer pair 515F-806R revealed higher number of taxa with increased abundance compared to other primer pairs. Among all the taxa showing significant differential abundance, the majority of species were from phyla Firmicutes and Actinobacteria. Archaeal taxa also showed higher abundance using the 515F-806R primer pair than when using other primer pairs (Figure 4).

**Figure 4.**
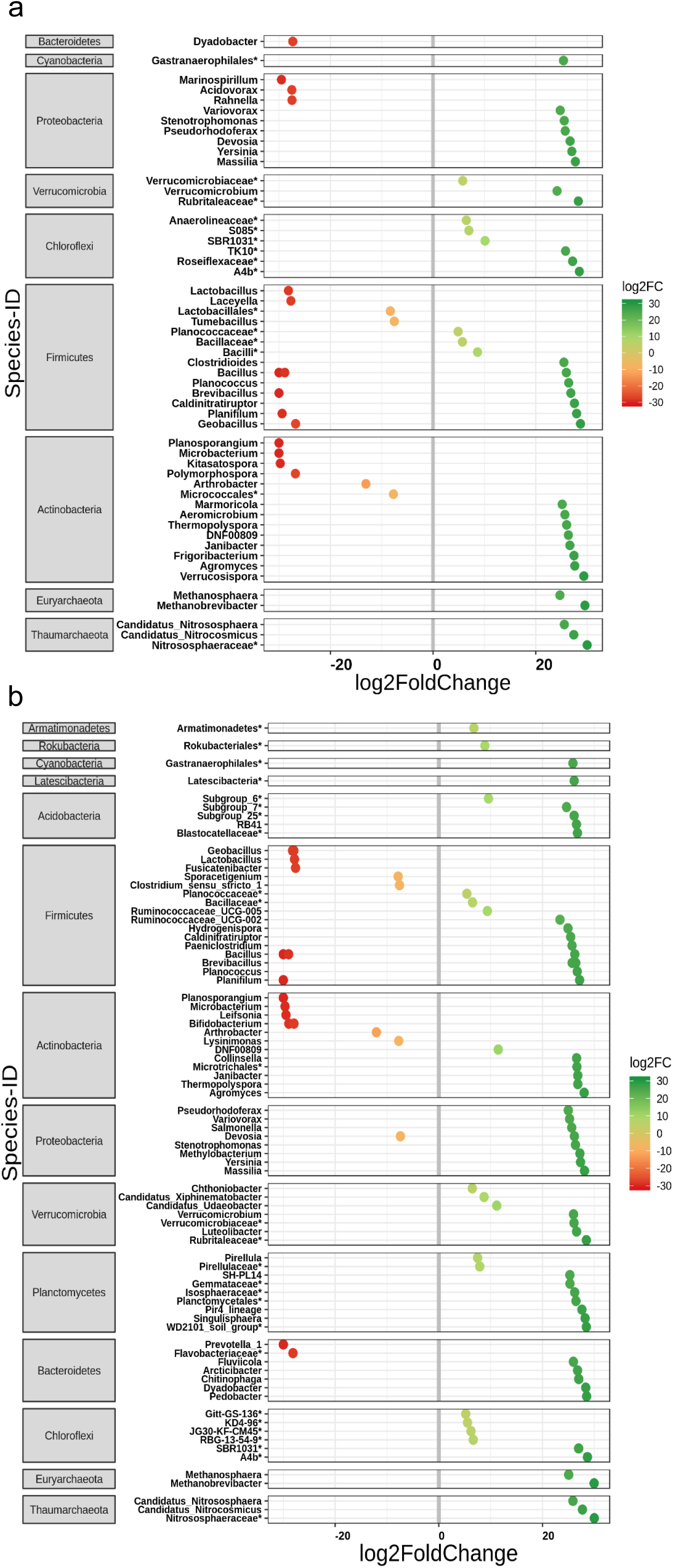
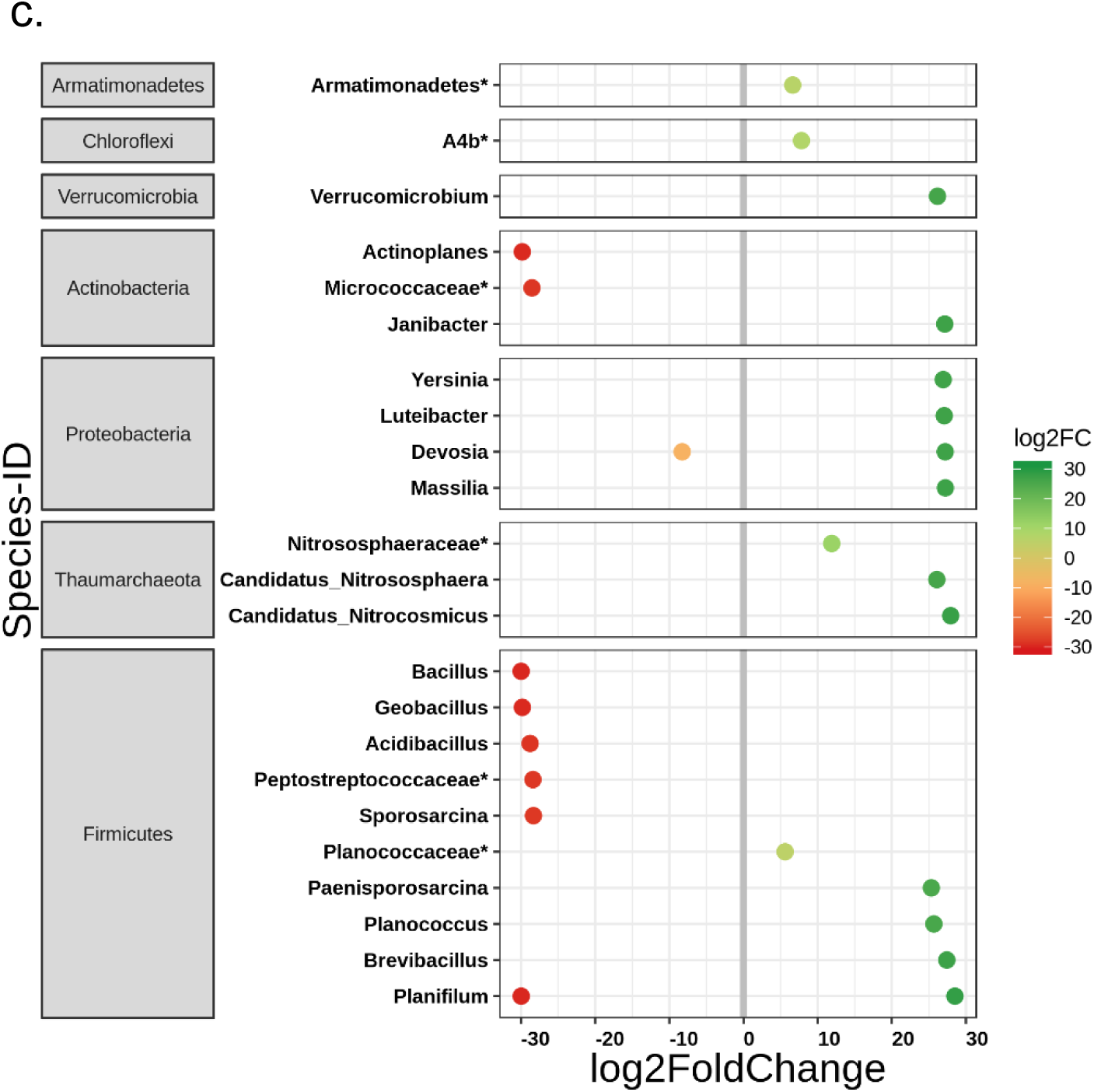
Differential abundance of species according to primer pair used. Shown are species that differ in their mean abundance in relation to the primer pair used. Primer pair 515F-806R was compared against **(a.)** 787F-1073R, **(b.)** 799F-1193R and **(c.)** 341F-805R. The values indicate a log2-fold (log2FC) decrease (red; using other primer pairs) or increase (green, using 515F-806R primer pair) in relative abundance of species. Species are arranged according to increasing values of log2-fold change and grouped according to their respective phylum. The highest possible taxonomic assignment (maximal to the genus level) is shown for each species. In all the comparisons, more species show increased abundance using 515F-806R than using any other primer pair.

### *In silico* primer pair evaluation

In order to evaluate the primer pairs *in silico*, weighted scores of primer matches were calculated for each primer pair against the latest SILVA v132 NR 16S rRNA gene database. Weighted score for individual primers was best (lowest) for 515F (0.09 ± 0.0006), followed by 341F (0.27 ± 0.0008), 806R (0.30 ± 0.0009), 1073R (0.43 ± 0.0012), 787F (0.51 ± 0.0015), 805R (0.62 ± 0.0019), 1193R (0.79 ± 0.0019) and 799F (0.89 ± 0.0022). Overall best score among the primer pairs was for 515F-806R (0.19), followed by 341F-805R (0.45), 787F-1073R (0.47) and 799F-1193R (0.84).

*In silico* primer coverage was predicted both for individual primers and pairs thereof at the domain (Archaea, Bacteria, Eukaryota) and phylum levels. At the domain level, primer coverage score was best for primer 515F, which covered all three domains (>90% coverage), whereas primer 1193R was found to be bacteria specific (Table 1). In terms of domain universality based on 80% coverage criteria, only primer 515F was found to be “universal” to all three domains, whereas 1073R and 1193R were bacteria specific. Likewise, both 515F-806R and 341F-805R were predicted universal for Bacteria and Archaea and two other tested primer pairs (787F-1073R, 799F-1193R) were found to be more bacteria specific (Table 1). Detailed coverage of primer pairs at domain and bacterial phylum levels are shown in Supplementary Table 6.

## Discussion

The inclusion of microbiomes as connecting links between trophic levels has been proposed in the One Health framework (van Bruggen et al., 2019). Before examining microbial transfers from one source compartment to another and implications for health at each level, methodological challenges for detecting and characterising microbiomes from trophic levels along the food chain must be met (Trinh et al., 2018). To overcome such limitations and to achieve a standard microbiome analysis approach, we investigated microbiomes from diverse source compartments along the food chain, using four commonly available primer pairs targeting different regions of 16S rRNA gene. As far as we know, this is the first integrated analysis of microbiomes along a food chain. We observed that among the tested primers all were explicitly targeting the bacterial domain and no reads belonging to Eukaryota were found. Primer pairs 515F-806R (P3) and 341F-805R (P4) performed better than others and also detected archaea as expected. Specifically, we recommend primer pair 515F-806R (P4) for One Health studies as it recovers the highest bacterial diversity both *in silico* and in samples from diverse source compartments along the food chain.

All the four tested primer pairs performed well and recovered the expected bacterial diversity in mock and mouse gut microbiomes. Nevertheless, we also noticed few rarer taxa in mock community samples (neglectable level of noise; <0.01%), which could be due to the fact that we did not perform taxa prevalence/abundance based filtering in our analyses because, each sample being unique in terms of microbiome composition in our study, such filtering would influence the four primer dataset differentially. As *DADA2* may be more sensitive to low amount of such contamination (Caruso et al., 2019), we recommend performing such filtering steps. We found that primer pair choice may significantly influence bacterial alpha diversity, which is supported by earlier studies that compared various primer pairs amplifying 16S rRNA gene (Beckers et al., 2016;Thijs et al., 2017). Highest microbial diversity was revealed by primer pair 515F-806R using two alpha diversity indices (number of observed species, Shannon). Our result is in contrast to previous studies (Klindworth et al., 2013;Thijs et al., 2017) where the primer pair 341F-805R showed the highest microbial diversity. Since the *in silico* study by *Klindworth et al*., the SILVA database nearly doubled in size following the inclusion of new sequences, and current primer coverage of the new database with existing primer pairs was up to now unknown. Meanwhile, there have also been several improvements made to the design of the forward and reverse primers of the 515F-806R pair in order to cover previously undetected taxa and to reduce biases against Crenarchaeota/Thaumarcheota (Apprill et al., 2015;Parada et al., 2016;Walters et al., 2016). Primer pair 515F-806R was not included in the comparative study by *Thijs et al*. because they intended to find additional suitable primer pairs for soil microbiome studies other than existing 515F-806R. Similar highest alpha diversity results for 515F-806R primer pair in comparison to other primer pairs were obtained in a recent study (Chen et al., 2019), however, limited to human gut microbiome. Primer pair 515F-806R has been the recommended primer pair for microbiome studies by the Earth Microbiome Project and has successfully been used in studies from other source compartments (Parada et al. 2016; Apprill et al. 2015; Walters et al. 2015), but, to the best of our knowledge, has never been tested with several compartments in the same study. We could not demonstrate a significant effect of extraction kit on overall alpha diversity, studies showing no effect or significant effect of DNA extraction kit on microbial alpha diversity are available in the literature (Hallmaier-Wacker et al., 2018;Fiedorova et al., 2019;Mattei et al., 2019).

The origin of the microbiota samples explained the majority of differences in alpha diversity patterns. Such differences in diversity were expected as each source compartment possesses its own microbial signature (Thompson et al., 2017;Ikeda-Ohtsubo et al., 2018;Reese and Dunn, 2018). Although, our study was focused on an agricultural food chain, the obtained alpha diversity results using 515F-806R primer pair are comparable with Earth Microbiome Project’s (EMP) findings, where similar primer pair was used to amplify diverse free-living and host-associated microbial communities (Thompson et al., 2017). We observed high microbial richness in free-living microbial community (i.e. in soil) compared to host-associated microbial communities (i.e. gut microbiome) similar to observations in the EMP. However, a notable exception was observed in the EMP, where plant roots showed highest microbial richness compared to all other host associated or free-living studied compartments (Thompson et al., 2017). We did not observe such patterns and among the studied source compartments, soil samples showed highest microbial richness in our study. Such discrepancy could arise due to differences in sample types (for example different species of plants), sampled root region (whole root processed in our experiment whereas the microbial rich rhizosphere was investigated in EMP) but also due to the fact that in EMP, root samples were collected from only two locations compare to worldwide equal distribution of collected soil samples, which showed a large microbial richness gradient (Thompson et al., 2017). In our study each source compartment exhibited unique microbiome, and only ∼17% of phyla were shared among microbial communities from different source compartments (soils, plant roots, cattle and humans) along the food chain. Four phyla; Firmicutes, Bacteroidetes, Actinobacteria and Proteobacteria dominate microbial communities along the model food chain, as each of them represent among major phyla in individual compartment specific studies (Hacquard et al., 2015;Ikeda-Ohtsubo et al., 2018). Furthermore, two archaeal phyla Euryarchaeota and Thaumarchaeota, only detected by primer pairs 515F-806R and 341F-805R, showed overall high abundance along the food chain. Species richness decreased gradually across the food chain, with the lowest richness observed in human faeces.

As observed for alpha diversity, beta diversity, as measured by Jaccard and Bray-Curtis dissimilarity indices, was significantly influenced by the choice of primer pairs; a finding that is supported by previous studies (Thijs et al., 2017;Chen et al., 2019). We observed no significant difference among primer pairs in beta-dispersion, suggesting that observed difference in beta diversity metrics among primer pairs are due to true difference in microbial community and not due to differential dispersion from centroids for each primer pair. As evidenced for alpha diversity, strong effect of source compartment on beta diversity is also expected as each source compartment harbours compositionally different sets of bacteria in different proportions (Hacquard et al., 2015;Thompson et al., 2017). Additionally, we found the factor interaction “primer pair*source compartment” to be significant in our model, thus indicating that some primer pairs may be better at revealing changes in microbial community composition than others for specific source compartment: For example, 515F-806R was associated with better spread of data points for soil samples in comparison to other primer pairs in the NMDS plot. Nevertheless, interpretation of beta diversity results is not as straightforward as for alpha diversity, because larger difference between samples, for example, can give overall high value for beta diversity, but this could arise due to limited, and thus differential, detection of taxa by a primer pair. Furthermore, no significant effect of extraction kit on beta diversity was observed in our analysis.

To investigate the observed differences in alpha and beta diversity using different primer pairs, we compared the relative abundances of bacterial species between the primer pair datasets. Overall, we found the largest number of differentially abundant taxa (86 species) when comparing primer pair 515F-806R with 799F-1193R, than comparing either 787F-1073R (58 species) or 341F-805R (25 species). Larger differences in case of 799F-1193R and 787F-1073R could be due to the fact that they amplify different hypervariable regions, V5-V6-V7 and V5-V6, respectively, as compared to the V4 region amplified by 515F-806R. Such discrepancies in abundance and taxa assignment of reads originating from different hypervariable regions of 16S rRNA gene have been reported previously (Claesson et al., 2010). In all comparisons performed in our analysis, higher proportion of bacterial species showed increased abundance when profiled with primer pair 515F-806R, thus reflecting the better performance of the latter primer pair over the three other primer pairs in detecting changes in taxon abundance. Many of the differential abundant species, which showed decrease or increase in all the comparisons belonged to Firmicutes and Actinobacteria, which are dominant phyla of microbiome community in different source compartments. Many species showing higher representation in 515F-806R primer pair in comparison with other primer pairs belonged to archaeal phyla, namely Euryarchaeota and Thaumarchaeota. Overall, a total of 78,659 (2.24%) reads were observed for these two phyla using primer pair 515F-806R as compared to 1,552 (0.09%) reads using the other prokaryotic primer pair 341F-805R. Euryarchaeota is a highly diverse archaeal phylum with predominating methanogenic species found from marine water to soil, plant root, cattle and human gut and it plays functionally important roles in each source compartment (Moissl-Eichinger et al., 2018). In humans, they are noted to be highly heritable and can be present in up to 95.7% of individuals in diverse studies (Lurie-Weinberger and Gophna, 2015). Their loss in ruminants can result in loss of energy and their role in obesity was suggested (Lurie-Weinberger and Gophna, 2015). Similarly, Thaumarchaeota commonly found in marine water, where they suggested to be keystone members of microbial community but also in diverse soil types and in association with plants (Brochier-Armanet et al., 2012;Taffner et al., 2019). Members of Thaumarchaeota are primarily known as ammonia oxidizers but also members with unknown energy metabolism (Ren et al., 2019). Considering the ubiquity and importance of Euryarchaeota and Thaumarchaeota in different source compartments, they could be important features in between-compartment studies and thus selection of prokaryotic primer pairs such as 515F-806R is highly recommended to amplify archaeal and bacterial taxa conjointly.

Our *in silico* analysis found 515F to be the most ‘universal’ among all tested primers, a finding similarly reported by *Klindworth et al.* Primer pair 515F-806R gave the best (lowest) weighted score followed by 341F-805R. In terms of coverage, we found both primer pairs 515F-806R and 341F-805R to be prokaryote specific with 515F-806R providing better coverage of archaea (96.39%) as compared to 341F-805R (83.59%), confirming our experimental observation, where higher proportion of reads assigned to archaea by primer pair 515F-806R than 341F-805R. However, primer pair 341F-805R was mentioned to be the best primer pair in terms of overall coverage by *Klindworth et al.* who used a smaller version of the SILVA database. Also, since then both the forward and reverse primers of the 515F-806R pair were made more general, i.e. wobble bases were added to both primers, and the reverse primer is now 6 base pair shorter in order to be less specific (Apprill et al., 2015;Parada et al., 2016;Walters et al., 2016).

In conclusion, we performed both laboratory and *in silico* tests to compare four commonly available 16S primer pairs in order to assess microbial community coverage across diverse source compartments along the food chain. Overall, we observed that the choice of primer pair can significantly influence microbial alpha and beta diversity and can identify differential taxa abundance. We showed that primer pair 515F-806R provides greater depth and taxa coverage as compared to other tested primer pairs using samples from different compartments, but also provides higher database coverage performance *in silico*. To conclude, we recommend including general prokaryotic primer pair such as 515F-806R to also recover archaea in One Health studies, due to their important roles in diverse systems. With this information, the methodological bottleneck concerning the choice of primer pair to adequately reflect microbial diversity within and between each source compartment can be addressed. This information will help identify and characterize the importance of microbiomes from heterogeneous origins within a One Health framework.

## Acknowledgements

This research was supported by the University of Bern through the Interfaculty Research Cooperation One Health (www.onehealth.unibe.ch). We are grateful to all the participating laboratories for providing source compartment specific samples. We thank the Interfaculty Bioinformatics Unit (IBU) Linux cluster from University of Bern and Institute for Infectious Diseases for providing computing facilities and Stefan Neuenschwander for assistance with software installation.

## Author’s contributions

ME, KS and AR were involved in the development of the conceptual framework for this study. AR and W conceived the study. KS and FR provided extracted DNA samples of soil/root and mouse, respectively. W carried out laboratory experiments, bioinformatics and statistical analyses. W wrote and AR edited the MS. All authors commented and approved the final version of the MS.

## Data accessibility

The individual microbial 16S rRNA gene sequences are available under BioProject ID PRJNA576426.

## Contribution to the field

Microbiomes not only provide essential services at each trophic level of the food chain i.e. soil, plant, cattle, human, but was recently recognised as a potential connecting links between trophic levels. Therefore, there is a need to track microbiome composition and transfer across diverse source compartments to understand microbial influence on the health of each compartment along the food chain. Yet before being able to compare compartments appropriately, methodological challenges to characterise microbial communities from diverse source compartments must be addressed. In that respect, we investigated the influence of different extraction kits and choice of 16S rRNA gene primer pairs on microbial community characterisation for the first time on diverse source compartments along the food chain. We observed that choice of primer pair can significantly influence microbiome characterisation. We make the recommendation of specific primer pair, 515F-806R which can cover highest microbial community diversity and providing best comparability between source compartments. Comparing microbiomes using 515F-806R revealed that soil and root samples have the highest estimates of species richness and inter-sample variation. Species richness decreased gradually along the food chain. Such knowledge has broader relevance, in making informed decisions concerning the choice of primer pair for microbiome researchers, irrespective of their studied source compartment and specifically can serve as a guide for those planning cross-compartment studies.

