## Supplementary Figures and Tables for "Evaluation of primer pairs for microbiome profiling across a food chain from soils to humans within the One Health framework"

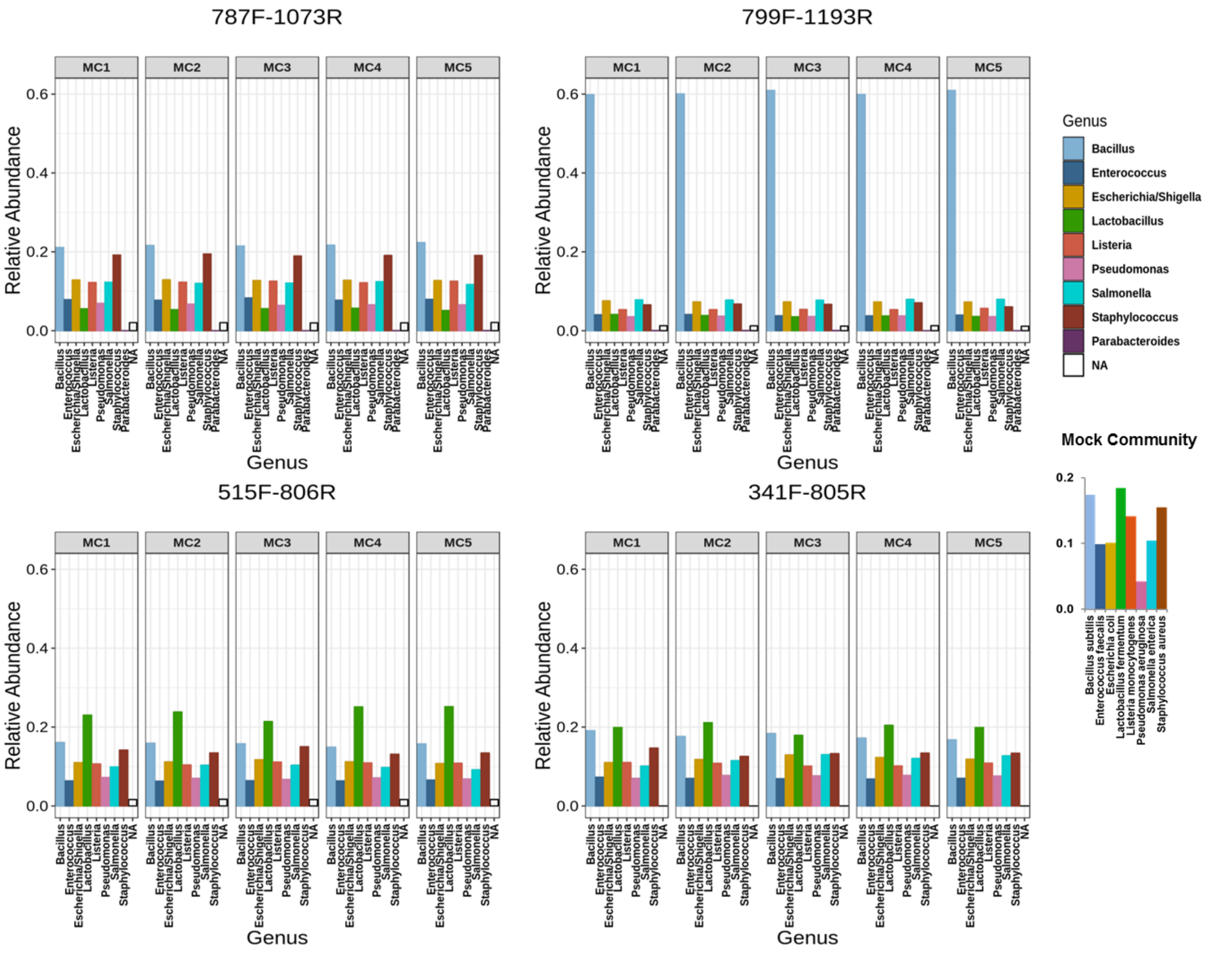

**Supplementary Figure 1.** Bar plots showing mock bacterial community at genus level as amplified by four (787F-1073R, 799F-1193R, 515F-806R and 341F-805R) primer pairs. In inset the expected proportions of bacterial species in mock community are shown. NA represents unassigned taxa at genus level.

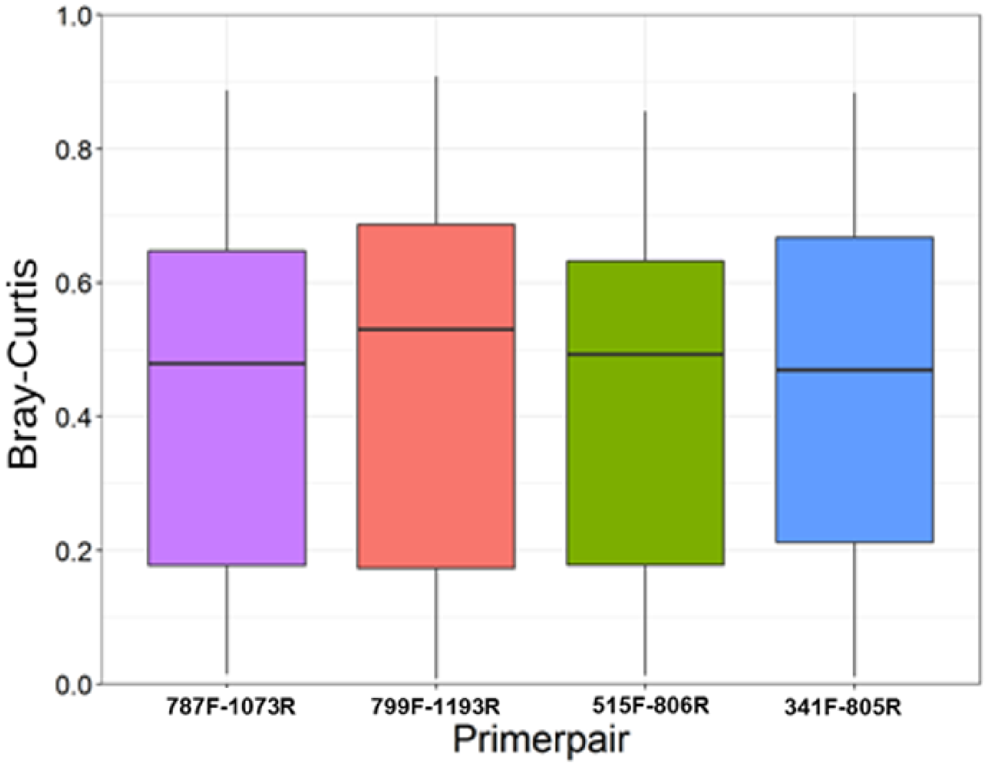

**Supplementary Figure 2.** Box plots showing average beta diversity based on Bray-Curtis metric for each prime pair. No significant difference in average beta diversity values was observed among primer pairs.

| Sample_ID | Source | Sample Characteristics | Sample Location |
|---|---|---|---|
| S1 | Soil | Soil-type1 | ARA_Uetendorf |
| S2 | Soil | Soil-type2 | Bodenacker |
| S3 | Soil | Soil-type3 | Kurzacker |
| S4 | Soil | Soil-type4 | Panorama |
| S5 | Soil | Soil-type5 | Q_matte |
| R1 | Maize-root | Maize-root-type1 | Ithaca |
| R2 | Maize-root | Maize-root-type2 | Changins |
| R3 | Maize-root | Maize-root-type2 | Changins |
| R4 | Maize-root | Maize-root-type2 | Changins |
| R5 | Maize-root | Maize-root-type3 | Reckenholz |
| R6 | Maize-root | Maize-root-type3 | Reckenholz |
| C1 | Cow | Animal1 | Vetsuisse |
| C2 | Cow | Animal2 | Vetsuisse |
| C3 | Cow | Animal3 | Vetsuisse |
| C4R | Cow-rumen | Animal4 | Vetsuisse |
| C5R | Cow-rumen | Animal5 | Vetsuisse |
| H1 | Human | Male_Age-44Y | Swiss-National |
| H2 | Human | Male_Age-02Y | Swiss-National |
| H3 | Human | Female_Age-26Y | Swiss-National |
| H4 | Human | Female_Age-43Y | Swiss-National |
| H5 | Human | Male_Age-38Y | Indian-National |
| H6 | Human | Female_Age-30Y | Indian-National |
| M1 | Mouse | Female_ASF4-colonized | DBMR |
| M2 | Mouse | Female_ASF4-colonized | DBMR |
| M3 | Mouse | Female_ASF4-colonized | DBMR |
| M4 | Mouse | Female_ASF4-colonized | DBMR |
| M5 | Mouse | Female_ASF4-colonized | DBMR |
| MC1 | Mock-community | Eight-bacterial-species | ZymoBIOMICS Microbial Community DNA Standard |
| MC2 | Mock-community | Eight-bacterial-species | ZymoBIOMICS Microbial Community DNA Standard |
| MC3 | Mock-community | Eight-bacterial-species | ZymoBIOMICS Microbial Community DNA Standard |
| MC4 | Mock-community | Eight-bacterial-species | ZymoBIOMICS Microbial Community DNA Standard |
| MC5 | Mock-community | Eight-bacterial-species | ZymoBIOMICS Microbial Community DNA Standard |

**Supplementary Table 1.** Details of the samples used in the current study.

| Primer pair | Raw Reads | QFiltered% | Denoised-F% | Denoised-R% | Merged% | Chimera% |
|---|---|---|---|---|---|---|
| 787F-1073R | 5,408,844 | 16.04 | 4.50 | 3.16 | 58.68 | 10.48 |
| 799F-1193R | 3,687,401 | 15.97 | 4.27 | 0.60 | 69.30 | 08.73 |
| 515F-806R | 4,898,357 | 10.61 | 1.66 | 0.80 | 80.90 | 04.15 |
| 341F-805R | 3,151,874 | 17.59 | 5.43 | 1.39 | 66.11 | 08.99 |

**Supplementary Table 2.** Distribution of reads at different stages of amplicon processing pipeline for used primer pairs.

| Primer pair | Bacteria | Chloroplast | Mitochondria | Archaea |
|---|---|---|---|---|
| 787F-1073R | 2,044,247 | 16 | 17,340 | 0 |
| 799F-1193R | 1,912,362 | 0 | 0 | 6 |
| 515F-806R | 3,507,030 | 41,597 | 61,908 | 79,066 |
| 341F-805R | 1,560,368 | 49,775 | 21,685 | 1552 |

**Supplementary Table 3.** Distribution of reads at domain level and for chloroplast and mitochondria for used primer pairs

| Variables | SumsOfSqs | MeanSqs | F.Model | p |
|---|---|---|---|---|
| ***(a) Observed species*** | | | | |
| Primerpair | 31604 | 10535 | 8.0362 | **<0.001** |
| Source compartment | 810819 | 162164 | 123.7055 | **<0.001** |
| Extraction kit | 49 | 49 | 0.0376 | 0.846 |
| Primer pair*Source compartment | 26790 | 1786 | 1.3624 | 0.171 |
| ***(b) Shannon*** | | | | |
| Primerpair | 4.848 | 1.616 | 11.2825 | **<0.001** |
| Source compartment | 181.966 | 36.393 | 254.0917 | **<0.001** |
| Extraction kit | 0.25 | 0.25 | 1.7454 | 0.188 |
| Primer pair*Source compartment | 3.805 | 0.254 | 1.771 | **0.042** |

**Supplementary Table 4.** Summary of the best model (GLM) predicting (a) Observed species and (b) Shannon diversity according to primer pair, source, extraction kit and primer pair*source compartment interaction. Significant values (<0.05) are shown in bold.

| Variables | SumsOfSqs | MeanSqs | F.Model | $R^2$ | p |
|---|---|---|---|---|---|
| ***(a) Jaccard*** | | | | | |
| Primerpair | 2.025 | 0.675 | 3.513 | 0.02464 | **0.001** |
| Source compartment | 39.965 | 7.9931 | 41.597 | 0.48633 | **0.001** |
| Extraction kit | 0.319 | 0.3188 | 1.659 | 0.00388 | 0.052 |
| Primer pair*Source compartment | 8.739 | 0.5826 | 3.032 | 0.10635 | **0.001** |
| Residuals | 31.129 | 0.1922 | | 0.3788 | |
| Total | 82.178 | | | 1 | |
| ***(b) Bray-Curtis*** | | | | | |
| Primerpair | 1.717 | 0.5722 | 4.401 | 0.02196 | **0.001** |
| Source compartment | 48.84 | 9.7679 | 75.124 | 0.62468 | **0.001** |
| Extraction kit | 0.192 | 0.1924 | 1.48 | 0.00246 | 0.142 |
| Primer pair*Source compartment | 6.371 | 0.4247 | 3.267 | 0.08149 | **0.001** |
| Residuals | 21.064 | 0.13 | | 0.26942 | |
| Total | 78.184 | | | 1 | |

**Supplementary Table 5.** Summary of the PERMANOVA models based on (a) Jaccard and (b) Bray-Curtis metric according to primer pair, source, extraction kit and primer pair*source compartment interaction. Significant values (<0.05) are shown in bold.

| Taxonomy level | Taxonomy for current level | Total seqs | 787F-1073R (P1) | 799F-1193R (P2) | 5151F-806R (P3) | 341F-805R (P4) |
|---|---|---|---|---|---|---|
| Archaea | Archaea | 25026 | 0.4914 | 0 | 0.9639 | 0.8359 |
| Bacteria | Bacteria | 592605 | 0.9691 | 0.8604 | 0.962 | 0.9669 |
| Eukaryota | Eukaryota | 77540 | 0 | 0 | 0.1663 | 0.0012 |
| Bacteria | Proteobacteria | 238929 | 0.9795 | 0.945 | 0.9799 | 0.9783 |
| Bacteria | Firmicutes | 149757 | 0.9743 | 0.9182 | 0.9662 | 0.964 |
| Bacteria | Actinobacteria | 60510 | 0.9799 | 0.9654 | 0.8796 | 0.9723 |
| Bacteria | Bacteroidetes | 55663 | 0.9669 | 0.9489 | 0.9742 | 0.9752 |
| Bacteria | Acidobacteria | 14534 | 0.9786 | 0.4586 | 0.9763 | 0.9788 |
| Bacteria | Cyanobacteria | 13970 | 0.8763 | 0.0088 | 0.9469 | 0.9465 |
| Bacteria | Chloroflexi | 9245 | 0.8852 | 0.2967 | 0.9447 | 0.9125 |
| Bacteria | Planctomycetes | 9014 | 0.9432 | 0.1295 | 0.9698 | 0.8776 |
| Bacteria | Epsilonbacteraeota | 5422 | 0.9572 | 0.8834 | 0.9817 | 0.9875 |
| Bacteria | Patescibacteria | 4521 | 0.7863 | 0.1694 | 0.8505 | 0.7436 |
| Bacteria | Verrucomicrobia | 4419 | 0.9446 | 0.1749 | 0.9615 | 0.9556 |
| Bacteria | Spirochaetes | 4253 | 0.925 | 0.858 | 0.902 | 0.9737 |
| Bacteria | Tenericutes | 2561 | 0.9559 | 0.4775 | 0.9805 | 0.977 |
| Bacteria | Fusobacteria | 2216 | 0.977 | 0.9666 | 0.9675 | 0.968 |
| Bacteria | Gemmatimonadetes | 2185 | 0.9611 | 0.9414 | 0.9593 | 0.9629 |
| Bacteria | Nitrospirae | 1297 | 0.9792 | 0.9545 | 0.9776 | 0.9807 |
| Bacteria | Synergistetes | 1152 | 0.9861 | 0.1615 | 0.9783 | 0.9583 |
| Bacteria | Kiritimatiellaeota | 975 | 0.96 | 0.799 | 0.9477 | 0.9631 |
| Bacteria | Deinococcus-thermus | 948 | 0.9631 | 0.9525 | 0.9789 | 0.9821 |
| Bacteria | Armatimonadetes | 752 | 0.9707 | 0.3191 | 0.9681 | 0.3218 |
| Bacteria | Fibrobacteres | 751 | 0.9587 | 0.9494 | 0.9614 | 0.9587 |
| Bacteria | Dependentiae | 580 | 0.981 | 0.8897 | 0.9897 | 0.9828 |
| Bacteria | Atribacteria | 578 | 0.9913 | 0.0969 | 0.9792 | 0.9723 |
| Bacteria | Marinimicrobia (sar406 clade) | 554 | 0.9838 | 0.9206 | 0.9819 | 0.9675 |
| Bacteria | Omnitrophicaeota | 507 | 0.9566 | 0.5089 | 0.9803 | 0.7456 |
| Bacteria | Latescibacteria | 497 | 0.994 | 0.4447 | 0.9819 | 0.9899 |
| Bacteria | Lentisphaerae | 469 | 0.9062 | 0.6269 | 0.9574 | 0.9659 |
| Bacteria | Chlamydiae | 450 | 0.9867 | 0.8178 | 0.9533 | 0.9889 |
| Bacteria | Elusimicrobia | 435 | 0.9724 | 0.9172 | 0.9885 | 0.9839 |
| Bacteria | Rokubacteria | 380 | 0.9737 | 0.0079 | 0.9711 | 0.9763 |
| Bacteria | Brc1 | 380 | 0.9684 | 0.0842 | 0.9421 | 0.9684 |
| Bacteria | Nitrospinae | 364 | 0.9698 | 0.022 | 0.9918 | 0.9863 |
| Bacteria | Aquificae | 346 | 0.9393 | 0.9595 | 0.9595 | 0.9855 |
| Bacteria | Thermotogae | 303 | 0.1716 | 0.769 | 0.9736 | 0.9868 |
| Bacteria | Cloacimonetes | 301 | 0.9668 | 0.8173 | 0.9435 | 0.9568 |
| Bacteria | Calditrichaeota | 275 | 0.9636 | 0.9564 | 0.9455 | 0.9418 |
| Bacteria | Hydrogenedentes | 271 | 0.9779 | 0.1771 | 0.9852 | 0.9668 |
| Bacteria | Halanaerobiaeota | 270 | 0.9667 | 0.0444 | 0.9667 | 0.9593 |
| Bacteria | Modulibacteria | 255 | 0.9686 | 0.0392 | 0.9255 | 0.9294 |
| Bacteria | Zixibacteria | 203 | 0.9901 | 0.9606 | 0.9655 | 0.9704 |
| Bacteria | Dadabacteria | 169 | 0.9704 | 0.8817 | 0.9763 | 0.9822 |
| Bacteria | Entotheonellaeota | 168 | 0.9286 | 0.9524 | 0.9702 | 0.9643 |
| Bacteria | Acetothermia | 165 | 0.9697 | 0.2364 | 0.9879 | 0.9939 |
| Bacteria | Fbp | 155 | 0.1935 | 0.8452 | 0.9484 | 0.9613 |
| Bacteria | Deferribacteres | 134 | 0.8881 | 0.9701 | 0.9925 | 0.9851 |

| | | | | | | |
|---|---|---|---|---|---|---|
| Bacteria | Gn01 | 120 | 0.9417 | 0.8 | 0.8833 | 0.0583 |
| Bacteria | Caldiserica | 95 | 0.8947 | 0.9263 | 0.9263 | 0.9684 |
| Bacteria | Ws1 | 93 | 0.9892 | 0.0323 | 0.9677 | 0.0323 |
| Bacteria | Pauc34f | 88 | 0.6364 | 0.7273 | 0.9773 | 0.9545 |
| Bacteria | Margulisbacteria | 86 | 1 | 0.4419 | 1 | 0.9535 |
| Bacteria | Coprothermobacteraeota | 76 | 0.8947 | 0.9474 | 0.9211 | 0.9474 |
| Bacteria | Ta06 | 68 | 0.8971 | 0.6765 | 0.9412 | 0.9853 |
| Bacteria | Aerophobetes | 66 | 0.9848 | 0 | 0.9091 | 0.9242 |
| Bacteria | Ws2 | 58 | 0.931 | 0.0172 | 0.9655 | 0.8621 |
| Bacteria | Aegiribacteria | 52 | 1 | 0.2692 | 0.9808 | 0.9808 |
| Bacteria | Wps-2 | 51 | 0.9804 | 0.0588 | 0.9412 | 0.9412 |
| Bacteria | Poribacteria | 49 | 0.9796 | 0.0204 | 0.9388 | 0.8776 |
| Bacteria | Lcp-89 | 46 | 0.9565 | 1 | 0.9783 | 1 |
| Bacteria | Gal15 | 43 | 1 | 0 | 1 | 0.9767 |
| Bacteria | Bhi80-139 | 41 | 0.9756 | 0.4146 | 0.9756 | 0.9756 |
| Bacteria | Schekmanbacteria | 40 | 0.975 | 0.025 | 1 | 1 |
| Bacteria | Hydrothermae | 38 | 0.9474 | 0 | 0.9737 | 0.9474 |
| Bacteria | Fcpu426 | 37 | 0.8919 | 0.2432 | 0.9189 | 0.8919 |
| Bacteria | Wor-1 | 34 | 0.9412 | 0.0882 | 0.9706 | 1 |
| Bacteria | Mat-cr-m4-b07 | 25 | 1 | 1 | 0.96 | 1 |
| Bacteria | Anck6 | 21 | 1 | 1 | 1 | 1 |
| Bacteria | Ck-2c2-2 | 21 | 0.9524 | 0.7619 | 1 | 1 |
| Bacteria | Ws4 | 16 | 1 | 0.0625 | 1 | 1 |
| Bacteria | Chrysiogenetes | 12 | 1 | 1 | 1 | 1 |
| Bacteria | Dictyoglomi | 11 | 1 | 0 | 1 | 1 |
| Bacteria | Rsahf231 | 8 | 1 | 0 | 1 | 1 |
| Bacteria | Uncultured | 5 | 1 | 0.2 | 1 | 1 |
| Bacteria | Fervidibacteria | 4 | 1 | 1 | 1 | 1 |
| Bacteria | Edwardsbacteria | 4 | 1 | 0.25 | 1 | 0.25 |
| Bacteria | Firestonebacteria | 3 | 1 | 1 | 1 | 1 |
| Bacteria | Calescamantes | 3 | 1 | 1 | 1 | 1 |
| Bacteria | Thermosulfidibacteraeota | 3 | 1 | 1 | 1 | 1 |
| Bacteria | Desantisbacteria | 2 | 1 | 1 | 1 | 1 |
| Bacteria | Gbs-1 | 2 | 1 | 1 | 1 | 1 |
| Bacteria | Lindowbacteria | 1 | 1 | 0 | 1 | 1 |

**Supplementary Table 6.** In silicocoverage (in proportion) of primer pairs at domain and phylum level.
